# Overcoming the impacts of two-step batch effect correction on gene expression estimation and inference

**DOI:** 10.1101/2021.01.24.428009

**Authors:** Tenglong Li, Yuqing Zhang, Prasad Patil, W. Evan Johnson

## Abstract

Non-ignorable technical variation is commonly observed across data from multiple experimental runs, platforms, or studies. These so-called batch effects can lead to difficulty in merging data from multiple sources, as they can severely bias the outcome of the analysis. Many groups have developed approaches for removing batch effects from data, usually by accommodating batch variables into the analysis (one-step correction) or by preprocessing the data prior to the formal or final analysis (two-step correction). One-step correction is often desirable due it its simplicity, but its flexibility is limited and it can be difficult to include batch variables uniformly when an analysis has multiple stages. Two-step correction allows for richer models of batch mean and variance. However, prior investigation has indicated that two-step correction can lead to incorrect statistical inference in downstream analysis. Generally speaking, two-step approaches introduce a correlation structure in the corrected data, which, if ignored, may lead to either exaggerated or diminished significance in downstream applications such as differential expression analysis. Here, we provide more intuitive and more formal evaluations of the impacts of two-step batch correction compared to existing literature. We demonstrate that the undesired impacts of two-step correction (exaggerated or diminished significance) depend on both the nature of the study design and the batch effects. We also provide strategies for overcoming these negative impacts in downstream analyses using the estimated correlation matrix of the corrected data. We compare the results of our proposed workflow with the results from other published one-step and two-step methods and show that our methods lead to more consistent false discovery controls and power of detection across a variety of batch effect scenarios. Software for our method is available through GitHub (https://github.com/jtleek/sva-devel) and will be available in future versions of the sva R package in the Bioconductor project (https://bioconductor.org/packages/release/bioc/html/sva.html). Batch effect; Two-step batch adjustment; ComBat; Sample correlation adjustment; Generalized least squares

## 1 Introduction

Because of the high cost of high-throughput profiling experiments or the difficulty in collecting a good number of samples, datasets are often processed in small batches, at different times, or in different facilities. These processing strategies often introduce unwanted technical variation into the data, commonly referred to as *batch effects*. RNA quality, lab protocol or experimenter, reagent batch, and other known and unknown factors affect the magnitude of batch effects and can often lead to significant technical heterogeneity and non-ignorable variation across batches (Leek and Storey, 2007, Leek et al., 2010, Johnson et al., 2007). It is well-established that batch effects will reduce statistical power and induce substantial bias for detecting differences between study groups (Leek et al., 2010, Johnson et al., 2007, Zhang et al., 2018). It is therefore common to perform some form of batch effect adjustment before the data are used for downstream analyses such as differential expression analysis (Leek and Storey, 2007).

There are many existing batch effect correction strategies, which can be classified as either “one-step” or “two-step” methods. One-step methods perform batch correction and data analysis simultaneously, by integrating the batch correction directly in the statistical model, prediction tool, or inference process. For example, a one-step strategy in a differential expression setting could be to include a batch indicator covariate in a linear model using common differential expression software tools (Smyth, 2005, Law et al., 2014, Robinson et al., 2010, Love et al., 2014). One-step approaches have the advantage of removing batch effects directly and succinctly in the modeling and analysis step. However, the batch correction is limited by the specific modeling approach, which in some cases may not adequately capture the behavior of the batch effects. In addition, one-step approaches may lead to inconsistent models or handling of the batch effects if multiple downstream steps are desired. For example, if a user wants to use hierarchical clustering for visualization, linear modeling for feature selection/identification, and non-linear machine learning (e.g. Support Vector Machines) for classification, it may be difficult to manage the batch effects for the entire analysis uniformly, because the batch adjustment will need to be incorporated into all steps in the same way.

In contrast, two-step methods perform batch correction as a data preprocessing step that is separate from the other steps of the analysis, outputting batch-corrected data for downstream tasks such as clustering, modeling, or prediction are applied to the data. There are several common methods for performing two-step batch correction, including ComBat (Johnson et al., 2007, Zhang et al., 2018, 2020), SVA (Leek et al., 2012), or RUV (Gagnon-Bartsch and Speed, 2012). Two-step methods such as ComBat are popular because they output “clean” data with batch effects removed, making the application of even complex downstream analyses more straightforward. Furthermore, adjusting for batch effects in a two-step process allows for the application of a richer model for batch adjustment (mean, variance, or other moments), which is often needed for combining highly heterogeneous batches of data or data from multiple studies.

The main drawback of two-step batch effect adjustment using methods such as ComBat is that it may lead to exaggerated significance if downstream modeling is not appropriately conducted, especially for unbalanced group-batch designs where the samples of a study group are distributed unevenly across batches (Nygaard et al., 2016). Consequently, the actual false positive rates (FPR) and false discovery rates (FDR) for some naïve downstream methods can be much higher than their nominal values, which renders results misleading. The root cause of exaggerated significance is the first step: removing batch effects with two-step methods (such as ComBat) introduces a correlation structure into the adjusted data. In a typical batch adjustment, the batch mean and/or variance are estimated using all the data points in the particular batch, and then this estimated batch mean is subtracted from each data point in the batch. This means that the adjusted data points within each batch are correlated with each other, because they are functions of all the other data from the batch. In addition to the exaggeration of significance established in previous work, we will show that in some circumstances this correlation structure can also result in diminished significance or power. Most researchers are unaware of these phenomena or are otherwise unable to incorporate this correlation structure into their models, which often leads to inappropriate downstream analyses that assume independent data points after batch correction.

In this article, we provide a basic theoretical explanation of the impacts of a naïve two-step batch correction strategy on downstream gene expression inference, and provide a heuristic demonstration and illustration of more complex scenarios using both simulated and real-data examples. We show that the group-batch design balance, i.e., whether the study/biological group design is correlated with the batch design, has a profound impact on the correlation structure induced by batch effect removal and thus on downstream analyses. We discuss the impact of the group-batch design balance on biological effect estimation and inference, and point out situations where we expect both exaggerated significance as well as diminished significance and power. We also propose a potential solution for mitigating the impacts of two-step batch correction on downstream analyses. Specifically, we show that the sample correlation matrix can be estimated for batch-corrected data and can be used in regression-based differential expression analysis (ComBat+Cor). This is equivalent to generalized least square estimation based on the estimated sample correlation matrix in batch-corrected data. The ComBat approach, combined with an appropriate variance estimation approach that is built on the group-batch design matrix, proves to be effective in addressing the exaggerated and/or diminished significance problem in ComBat-adjusted data.

## 2 Methods

### 2.1 Two-step batch adjustment and sample correlation

To illustrate the correlation structure introduced by two-step batch adjustment methods, we describe a simplified problem with a mean/additive batch effect only. Based on ComBat (Johnson et al., 2007), we describe gene expression data with only batch effects in the mean with the following linear model:

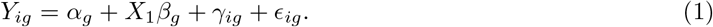

*Y*_*ig*_ denotes the gene expression of gene *g* for samples from batch *i*, which is the sum of the background expression *α*_*g*_, the vector of biological effects *β*_*g*_ corresponding to a biological group design matrix *X*_1_, the mean batch effects *γ*_*ig*_ for batch *i* and the residual term *∈*_*ig*_. Without loss of generality, we reformulate the above equation in matrix form:

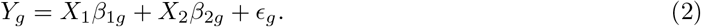

For the model above, we define *X* = [*X*_1_, *X*_2_] such that the matrix *X*_1_ consists of the indicators of biological groups (the group design) and the matrix *X*_2_ consists of the indicators of batches (the batch design). Therefore, *X* represents the group-batch design and is of central interest in this paper. In this case we will assume the errors *∈*_*g*_ follows a Normal distribution 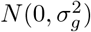. The model (2) can be used to adjust for mean batch effects and we refer to this approach as the “one-step” approach. Alternatively, batch effect adjustment can be done by a two-step approach. In the first step, the batch effects are estimated by 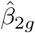 based on the regression (2) above and the batch-adjusted data 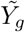 is obtained by removing the estimated batch effects from *Y*_*g*_, i.e., 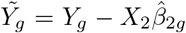 The variance of the adjusted data 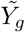 can be estimated as 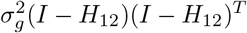, where 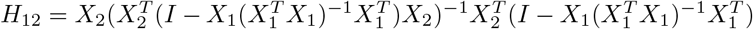. For a reference batch (Zhang et al., 2018), the matrix *X*_1_ should also include the all-ones vector **1** (see the supplementary material for derivation).

In the second step of the two-step approach using a similar linear modeling approach, the biological effect *β*_1*g*_ is estimated based on the adjusted values 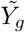:

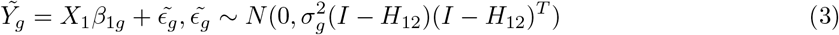

As derived above (and in the supplementary material), the samples in adjusted data 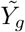 are correlated, with the correlation matrix defined by *M* = (*I* − *H*_12_)(*I* − *H*_12_)^*T*^. One interesting result is that in balanced group-batch designs, i.e., samples of a biological group are uniformly distributed across batches, the correlations of the adjusted data values are only dependent on the batch design, and not the group design (see Supplementary Methods, Section 3 for derivation). However, in unbalanced group-batch designs, the correlations among individual adjusted values, both within and across batches, depend also on the group design, which may have an influential impact on downstream analysis. Regardless of whether or not the group-batch design is balanced, researchers need to apply downstream analyses that are appropriate for correlated data, as these correlations may in some cases have profound impact on statistical inference of the biological effects if not properly modeled.

### 2.2 Impact of the design balance on biological effect estimation

One important implication of the covariance structure defined above is that the correlation matrix *M* may depend on the biological group design *X*_1_, resulting in possible correlations between the residuals and the covariate itself in (3), a concept often termed as *endogeneity*. In this section, we will show that this issue is related to the balance of the group-batch design, i.e., whether or not the group design is correlated with the batch design. To start, we derive the formula for 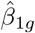 in the model (2) as (see the supplementary material):

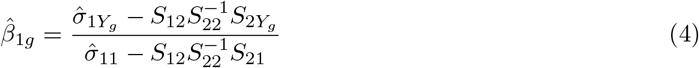

where *S*_12_ is the covariance matrix between the group design *X*_1_ and the batch design *X*_2_, *S*_22_ is the covariance matrix of *X*_2_, 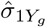 is the sample covariance of *X*_1_ and the outcome *Y*_*g*_, and 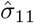 is the sample variance of *X*_1_. The critical piece in (4) from an endogeneity perspective is *S*_12_. We will now consider cases where the batch-covariate design is balanced and unbalanced.

#### 2.2.1 Balanced designs

If the group-batch design is balanced, *S*_12_ = 0, and the expression in (4) can be simplified to 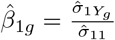. Thus it is only important to accurately estimate the residual variance using the adjusted data. We note that this still requires knowledge of the batch design *X*_2_. However, in gene expression analysis, the correlation structure for balanced designs is the same for all genes, providing ample data to estimate the correlation structure even if the overall sample size is small. It follows that the biological effect estimates 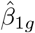 are the same for the following three models:

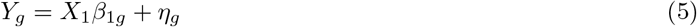

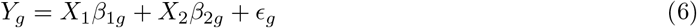

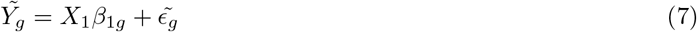

This suggests that the biological effect estimate 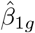 is not affected by batch effect and therefore the *endogeneity* issue does not exist in balanced batch-group design. This is because for balanced batch-group designs the adjusted data correlations do not depend on the group design *X*_1_. We make two important observations here: first, the variance for 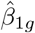 in the first model will be larger than the variance in the second model, especially if the batch effect is significant. This means that excluding the batch effect term from the model will not bias the estimate of the biological effect, but it will inflate the estimate for the residual standard deviation, leading to a reduction in power. Second, the variance of 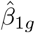 in the third equation can be estimated using ordinary least squares with the appropriate mean squared error estimate, or using generalized least squares to directly estimate the residual variance across all genes.

#### 2.2.2 Endogeneity in unbalanced designs

In unbalanced design, the expressions in (3) and (4) have clear implications in batch correction contexts that must be considered carefully. First, because the columns of the group design *X*_1_ and the batch design *X*_2_ are not linearly independent, and the *S*_12_ covariance matrix is non-zero, unlike the balanced design case. Second, the adjusted data correlations are dependent on both the batch and group designs, and the correlation structure will depend on the nature and magnitude of the biological effects. Therefore, this correlation structure will be different across the genes and cannot be easily estimated in gene expression data with small sample sizes.

### 2.3 Exaggerated and diminished significance in differential expression analysis

The endogeneity in unbalanced designs can bias the biological effect and variance estimates in gene expression analysis, often leading to incorrect *p*-values for downstream differential expression. In general, the correlation structure induced by two-step adjustments leads to the underestimation of the residual error if correlation is ignored. As a result, the two-step approach for the model in (1) usually results in artificially smaller *p*-values, and inflates the false positive rate (FPR) or false discovery rate (FDR) if the correlation not properly modeled. Typically, the level of FPR inflation increases as the group-batch design becomes more unbalanced. This phenomenon is often referred to as *exaggerated significance* (Nygaard et al., 2016). To overcome the exaggerated significance problem, the correlation matrix *M* needs to be computed and accounted for in the two-step analyses.

Some batch correction methods, such as ComBat, use a richer model than that of (1), in that they model and correct for both mean batch effects *γ*_*ig*_, and variance batch effects *δ*_*ig*_:

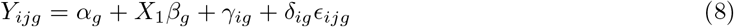

If variance batch effects are not present or negligible, i.e., *δ*_*ig*_ are all close to 1, the model in (8) is equivalent to the mean-only batch model in (1), for which we have derived the correlation matrix *M* for batch-corrected data. In this case, using ComBat for batch correction will also result in exaggerated significance for unbalanced designs, as previously described, if methods for correlated data or the correlation matrix *M* are not used in downstream modeling.

However, if variance batch effects are large, i.e., *δ*_*ig*_ are significantly different among batches and genes, failure to consider the correlation can have very different effects, possibly leading to either exaggerated or diminished significance. Intuitively, one factor that could drive this phenomenon is an underestimation or overestimation of the residual variance. Specifically, for the ComBat model, the residual variance estimate is given by the following (Johnson et al., 2007):

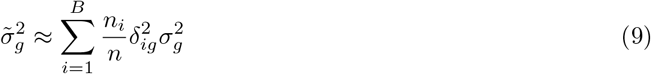

where 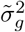 is the ComBat estimate for the residual variance, *n*_*i*_ is the size of *i*^*th*^ batch, and the total sample size is *n*. For identifiability, the ComBat model calibrates the *δ*_*ig*_ so that their products are equal to 1. The variance estimate 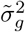 may be smaller than expected if some or all *δ*_*ig*_ are significantly less than 1 due to estimation error. In this case, this underestimation would lead to exaggerated significance. On the other hand, 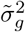 would be overestimated if most of the *δ*_*ig*_ are larger than 1, which likely leads to a conservative FPR and potentially a significant loss of statistical power. Therefore, variance batch effects mainly affect the estimate of residual variance 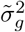, as evidenced by the contrast between (1) and (8), and large variance batch effects likely lead to a different and much more complicated expression of *M*. For the inference of biological effect, this means the statistical significance can be either exaggerated or diminished using ComBat, depending on the distributions of 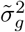 among batches and genes. We caution readers to be specific about variance batch effects when discussing the exaggerated significance problem for ComBat.

### 2.4 Computing the correlation matrix

We propose additional steps to appropriately address the correlation structure in two-step adjusted data: First, the sample correlation matrix introduced by batch effect removal needs to be estimated. Then, any downstream analysis based on batch-corrected data needs to utilize the sample correlation matrix in their correlated data models. Obtaining *M* = (*I* − *H*_12_)(*I* − *H*_12_)^*T*^ is straightforward given both the batch and biological design matrices, *X* = [*X*_1_, *X*_2_], except that it may not be not full rank due to the batch correction. Thus *M* needs to be approximated by another full rank matrix 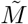 in order to make it usable for downstream analysis. We note that *M* is not gene-specific and does not consider the differences among gene-specific covariance matrices in unbalanced group-batch design.

We propose to use the following steps to obtain an approximated sample correlation matrix 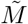: (1) Apply a spectral decomposition *M* = *Q*Λ*Q*^*T*^, where *Q* consists of the of eigenvectors of *M* and Λ is the diagonal matrix with eigenvalues of *M* as its diagonal elements; (2) Since the batch-corrected data is obtained by removing mean batch effect estimates from every observation, *M* is not full rank and has some zero eigenvalues. We will replace those zero eigenvalues by a small non-zero number, *θ* (Cheng and Higham, 1998, Zusmanovich, 2013). Conceptually, this is equivalent to adding a small amount of random noise to the dataset to make it full rank. We will denote the modified set of eigenvalues, with zeros replaced by *θ*, as 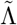; (3) The approximated sample correlation matrix is computed by 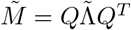. To enhance interpretability, we will redefine *θ* as the product of the sum of non-zero eigenvalues and *ζ*, in which *ζ* represents the percentage of noise added by the user. It is recommended that *ζ* should be chosen as a value between 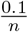 and 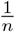 (Knol and ten Berge, 1989), where *n* is the total sample size of the combined batches, as both under-adjustment and over-adjustment may negatively influence statistical power. We will demonstrate the impact of *ζ* on batch effect adjustment using our simulation studies below.

### 2.5 Use the sample correlation matrix in differential expression analysis

Based on 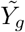 and 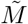, the linear model for differential expression analysis becomes:

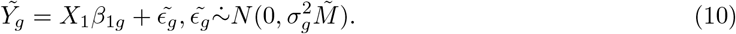

The biological group effects can then be estimated through methods for correlated data, such as generalized least squares (GLS). This will require a Cholesky decomposition of 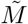. Linear transformation of both 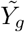 and *X*_1_ based on the Cholesky decomposition are also required but will be straightforward. Given that 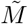 informs the sample correlations brought by the unbalanced design, the batch-corrected data is no longer correlated via the transformation based on 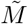 and therefore GLS estimation should lead to proper statistical significance.

Here we propose an enchanced version of ComBat, ComBat+Cor (the ComBat approach that includes a correlation adjustment). Incorporating the above procedure will mitigate downstream impacts such as exaggerated *p*-values (and *q* - values) for unbalanced group-batch designs. ComBat+Cor comprises the following three steps:

1. Use the original ComBat approach to obtain batch-adjusted data.
2. Obtain the sample correlation matrix 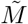 based on the design matrix *X* and the noise parameter *ζ*.
3. Use downstream analysis methods that accommodate correlated data. For example, estimate the group effect(s) and variance estimates using GLS based on 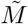.

### 2.6 Simulation design

To illustrate the effectiveness of ComBat+Cor in addressing the exaggerated significance problem for unbalanced designs, we first performed experiments on simulated data with batch effects. Datasets were simulated with mean and variance batch effects at different levels in order to examine the impact of batch effect sizes on the effectiveness of ComBat+Cor.

In our experiments, we simulated datasets based on the experimental design of a previously evaluated bladder cancer dataset, henceforth denoted as the *bladderbatch* data (Dyrskjøt et al., 2004, Leek et al., 2010). The simulated bladderbatch datasets followed the original study design for batches and cancer status, which was highly unbalanced with respect to status and batch. There were five batches in total, and the numbers of cancer/control samples in each batch were 11/0, 14/4, 0/4, 0/5, and 15/4, respectively. For comparison, we also simulated datasets based on a balanced group-batch design. The number of treated/control samples in the balanced design in each batch were 6/6, 9/9, 2/2, 3/3, and 10/10, respectively. We simulated the expression of 20,000 genes, of which 2,000 were set to be differentially expressed between the groups (treatment versus control). Group effects for the 2,000 differentially expressed genes were chosen as 2 (500 genes), −1 (500 genes), −1 (500 genes), −2 (500 genes), reflecting scenarios when the group effect was strongly positive, positive, negative and strongly negative. The remaining 18,000 genes were not differentially expressed between biological groups (“null” genes).

Our simulation method follows the hierarchical linear model assumed in ComBat given in Equation 8 (Johnson et al., 2007). We also specified the number of samples, batches, and genes in the data, and the distributions of mean and variance batch effects are given as *γ*_*ig*_ ~ *N* (*m*_*i*_, *v*_*i*_) and *δ*_*ig*_ ~ IG(*α*_*i*_, *β*_*i*_) for batch *i*. We then sampled *γ*_*ig*_ and *δ*_*ig*_ from their hyperparameter distributions. The background average expression *α*_*g*_ was set to be 3, and the gene-wise variation followed a gamma distribution Γ(4.5, 1.5). For the residuals *∈*_*ijg*_, the variances 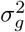 were randomly drawn from a gamma distribution Γ(4, 10) and we randomly sampled *∈*_*ijg*_ from 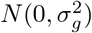. The above parameters were chosen based on the ComBat estimates for the original *bladderbatch* data, and were later modified to reflect scenarios where the batch effects were much larger than the original estimates (see Table 1). With simulated batch effects, the final gene expression *Y*_*ijg*_ was calculated as *Y*_*ijg*_ = *α*_*g*_ + *X*_1_*β*_*g*_ + *γ*_*ig*_ + *δ*_*ig*_*∈*_*ijg*_. To set a benchmark for simulation, we generated data without batch effect as 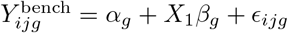.

**Table 1:** Hyperparameters of the mean and variance batch effects used in the simulation studies. The “Data” column refers to the parameter values estimated based on the original data. The “Small” and “Large” columns refer to the parameter values used for simulating data with small and large batch effects respectively.

| Batch | $m_i$ | | | $v_i$ | | | $\alpha_i$ | | | $\beta_i$ | | |
| --- | --- | --- | --- | --- | --- | --- | --- | --- | --- | --- | --- | --- |
|  | Data | Small | Large | Data | Small | Large | Data | Small | Large | Data | Small | Large |
| 1 | -0.04 | -0.04 | -0.4 | 0.15 | 0.15 | 0.15 | 60 | 60 | 100 | 60 | 60 | 100 |
| 2 | 0.15 | 0.15 | 1.5 | 0.35 | 0.35 | 0.35 | 100 | 100 | 120 | 100 | 100 | 40 |
| 3 | -0.15 | -0.15 | -1.5 | 0.82 | 0.82 | 0.82 | 56 | 56 | 100 | 50 | 50 | 60 |
| 4 | -0.1 | -0.1 | 1.0 | 0.46 | 0.46 | 0.46 | 30 | 30 | 60 | 30 | 30 | 100 |
| 5 | -0.08 | -0.08 | -0.8 | 0.12 | 0.12 | 0.12 | 100 | 100 | 40 | 100 | 100 | 120 |

We ran a differential expression analysis using a linear model on data without batch effects, and used the *p*/*q* - values obtained in this approach as the benchmark for the uncorrected and batch-corrected data. After including the batch effects, we compared ComBat and ComBat+Cor in terms of their distributions of *p*-values and false discovery rates. ComBat+Cor, as mentioned earlier, relies on the value of the noise parameter *θ*. Therefore, we ran ComBat+Cor with different values of *θ* to check the sensitivity of ComBat+Cor with regard to the choice of *θ*. T-tests based on the unadjusted raw data were conducted in order to illustrate the necessity of batch effect adjustment. Results based on the one-step approach, which controls for both the group and batch indicators in regression model, were also included.

### 2.7 Empirical examples

In addition to the simulation study based on the *bladderbatch* data, we provide three real data examples that have unbalanced group-batch designs. The first example is a dataset from Towfic et al. [2014], which is used to compare the effects of Copaxone and Glatimer. The second example is a dataset from Johnson et al. [2007], which is used for comparison of TAL1 inhibited cells. The first and second examples were actually used by Nygaard et al. [2016] to illustrate the exaggerated significance problem in ComBat. The third example is from several tuberculosis (TB) gene expression studies (Zak et al., 2016, Suliman et al., 2018, Leong et al., 2018), and we compare the gene expressions of progressors versus non-progressors in TB. For each example, we compare the *p*/*q* - values of ComBat and ComBat+Cor. We also conduct simulations based on mean and variance batch effects estimated by ComBat for all three examples, and for each simulation we compare the *p*-values based on the benchmark approach (dataset without batch effects), ComBat and ComBat+Cor, to illustrate the effectiveness of ComBat+Cor in these examples.

## 3 Results

We provide results for our primary *bladderbatch* simulation experiment (Example 1), re-analyses of two examples described by Nygaard et al. [2016] (Examples 2 and 3), and an additional case in tuberculosis gene expression with pronounced variance batch effects (Example 4).

### 3.1 Example 1: Simulated bladderbatch datasets

For datasets simulated based on the original bladderbatch data, we found that ComBat generated exaggerated *p*-values compared to the benchmark *p*-values (Figure 1(a)). The false positive rate (FPR) for ComBat was 18.3% which was much higher than the nominal rate of 5% (Table 2). In contrast, ComBat+Cor (with *ζ* = 1%) was able to appropriately control the false positive rate (Figure 1(b)). The FPR for ComBat+Cor (with *ζ* = 1%) was 4.8%. The distributions of *p*-values for the benchmark, ComBat and ComBat+Cor are depicted in Figure 1(c). Unsurprisingly, ComBat also yielded an exaggerated FDR that was much higher than the nominal one. We identified 3,264 genes as differentially expressed using ComBat and an FDR = 5% as the threshold. Of these genes, 1,978 were truly differentially expressed, yielding a detection power of 98.9%. The remaining 1,286 were actually “null” genes (i.e. genes not differentially expressed), meaning the actual FDR was inflated to 39.4%. Using ComBat+Cor with *ζ* = 1%, 2,001 genes were identified as significant using the same FDR cutoff, 1,926 of which were truly differentially expressed (power=96.3%). Only 75 of the genes identified as differentially expressed were null genes, yielding an actual FDR for ComBat+Cor of 3.7%. Therefore, the ComBat+Cor method provided considerably improved FDR control while retaining high detection power as compared to ComBat.

**Figure 1:**
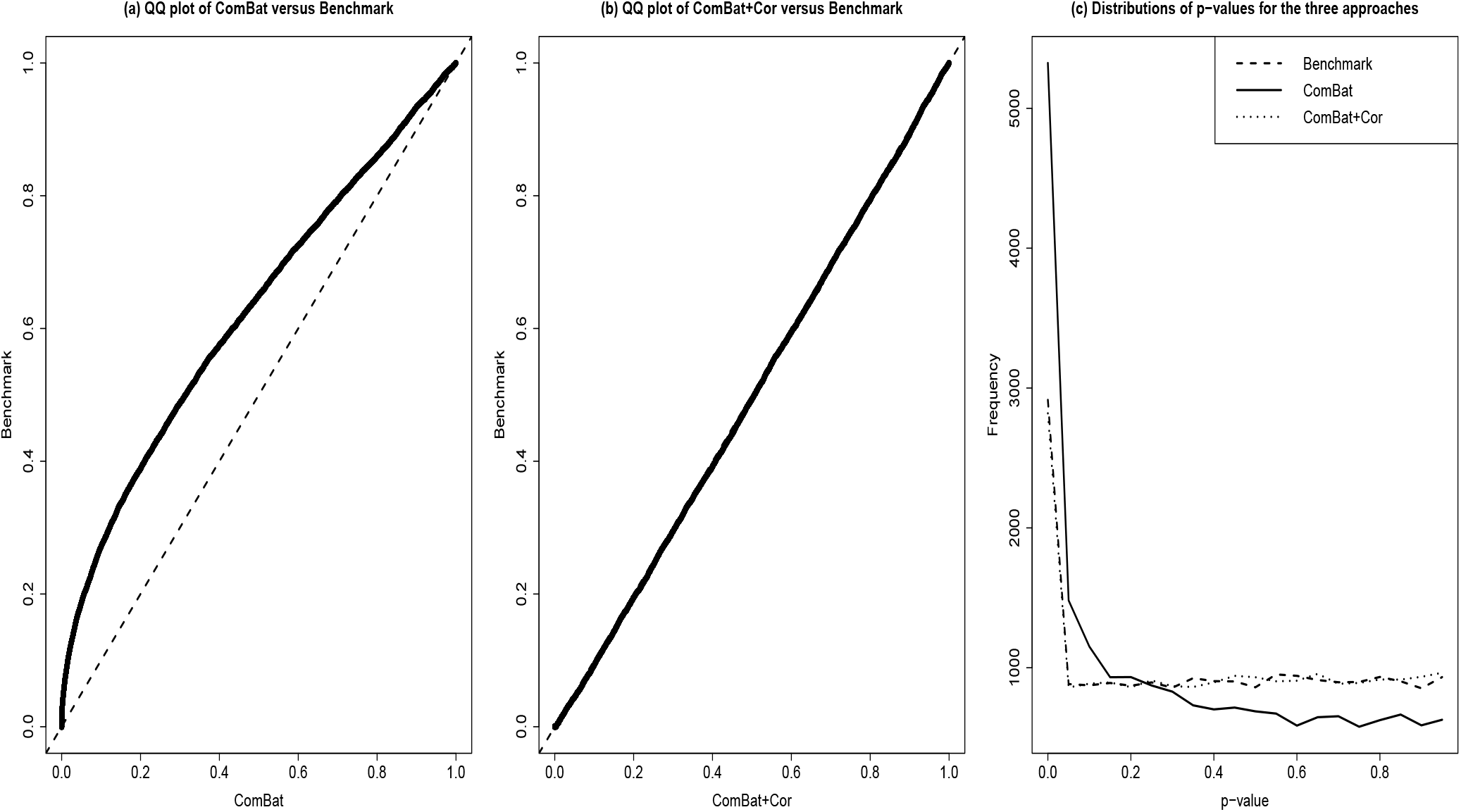
Three figures are used to illustrate that ComBat+Cor reduces the exaggerated significance seen when ComBat is applied based on simulated data that mimics the *bladderbatch* experimental design. Note that the original *bladderbatch* data has unbalanced group-batch design and small (mean and variance) batch effects. The benchmark approach refers to the approach that applies ordinary differential expression analysis to data without any batch effects. (a) QQ plot of *p*-values using ComBat and the *p*-values using the benchmark approach. The line falls above the *y* = *x* identity line, suggesting that *p*-values generated by ComBat concentrate at smaller values than those generated on the data without batch effect. (b) QQ plot of *p*-values using ComBat+Cor (*ζ* = 1%) and *p*-values using the benchmark approach. (c) line chart comparing the distributions of *p*-values using ComBat, ComBat+Cor and the benchmark approach.

**Table 2:** Results from *bladderbatch* simulation with unbalanced design. For each approach, results were obtained under the conditions where the mean and variance batch effects could be null (N), small (S) or large (L). For each condition, the results are formatted as FPR (TPR).

| Approach | Mean(N) |  |  | Mean(S) |  |  | Mean(L) |  |  |
| --- | --- | --- | --- | --- | --- | --- | --- | --- | --- |
|  | Var(N) | Var(S) | Var(L) | Var(N) | Var(S) | Var(L) | Var(N) | Var(S) | Var(L) |
| T-test | 4.9% | 4.8% | 4.2% | 34.0% | 33.3% | 22.6% | 37.8% | 38.0% | 31.2% |
|  | (99.7%) | (99.9%) | (95.9%) | (97.8%) | (97.3%) | (93.1%) | (78.9%) | (79.1%) | (76.8%) |
| Benchmark | 4.9% | 4.8% | 5.3% | 5.3% | 5.0% | 5.1% | 4.8% | 5.0% | 5.0% |
|  | (99.7%) | (99.9%) | (99.8%) | (99.7%) | (99.9%) | (99.9%) | (99.9%) | (99.7%) | (99.8%) |
| One-step | 4.9% | 5.0% | 8.4% | 5.2% | 5.1% | 8.1% | 4.9% | 5.0% | 8.3% |
|  | (99.1%) | (99.0%) | (89.0%) | (98.8%) | (98.5%) | (90.0%) | (98.7%) | (99.1%) | (89.7%) |
| ComBat | 5.6% | 10.9% | 1.0% | 14.6% | 18.3% | 3.4% | 15.5% | 18.5% | 3.5% |
|  | (99.7%) | (99.9%) | (98.4%) | (99.7%) | (99.6%) | (98.2%) | (99.6%) | (99.6%) | (98.2%) |
| ComBat+Cor($\zeta = 10\%$) | 0.0% | 0.0% | 0.0% | 0.2% | 0.2% | 0.0% | 0.2% | 0.3% | 0.0% |
|  | (94.8%) | (96.1%) | (78.8%) | (94.9%) | (95.7%) | (78.1%) | (94.7%) | (95.3%) | (78.4%) |
| ComBat+Cor($\zeta = 1\%$) | 0.4% | 1.6% | 0.2% | 2.7% | 4.8% | 0.6% | 3.0% | 4.7% | 0.6% |
|  | (99.0%) | (99.2%) | (93.2%) | (98.7%) | (98.6%) | (94.0%) | (98.6%) | (99.1%) | (93.4%) |
| ComBat+Cor($\zeta = 0.1\%$) | 0.2% | 1.1% | 0.1% | 1.3% | 6.3% | 0.3% | 2.1% | 6.6% | 0.3% |
|  | (97.4%) | (99.1%) | (92.1%) | (98.3%) | (98.8%) | (92.4%) | (98.2%) | (99.4%) | (92.1%) |
| ComBat+Cor($\zeta = 0.001\%$) | 0.0% | 0.0% | 0.0% | 0.0% | 1.3% | 0.0% | 0.0% | 2.5% | 0.0% |
|  | (23.8%) | (51.3%) | (21.7%) | (17.8%) | (97.9%) | (23.0%) | (33.6%) | (98.8%) | (28.8%) |

In addition, we conducted a sensitivity analysis using multiple values of *ζ*. Our earlier recommendations for *ζ* (between 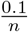 and 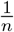 for sample size *n*) yield values in the [0.17%, 1.7%] range for the sample size of this study (*n* = 57). We ran ComBat+Cor with different values of *ζ* in order to evaluate the recommended range, as well as check the sensitivity of ComBat+Cor to values outside the range. Figure 2 presents the true positive rate (TPR) associated with different *ζ* values. These results suggest that when *ζ* is smaller than 2% and larger than 0.1% (consistent with the recommended range), ComBat+Cor achieved acceptable power (> 95% for both unbalanced and balanced designs) for detecting differentially expressed genes. ComBat+Cor lost power when *ζ* was either above or below the recommended range. Meanwhile, the FPR was consistently below 5% across this range of *ζ* values, signaling that Combat+Cor will produce conservative results regardless of choice of *ζ*. (Figure 3).

**Figure 2:**
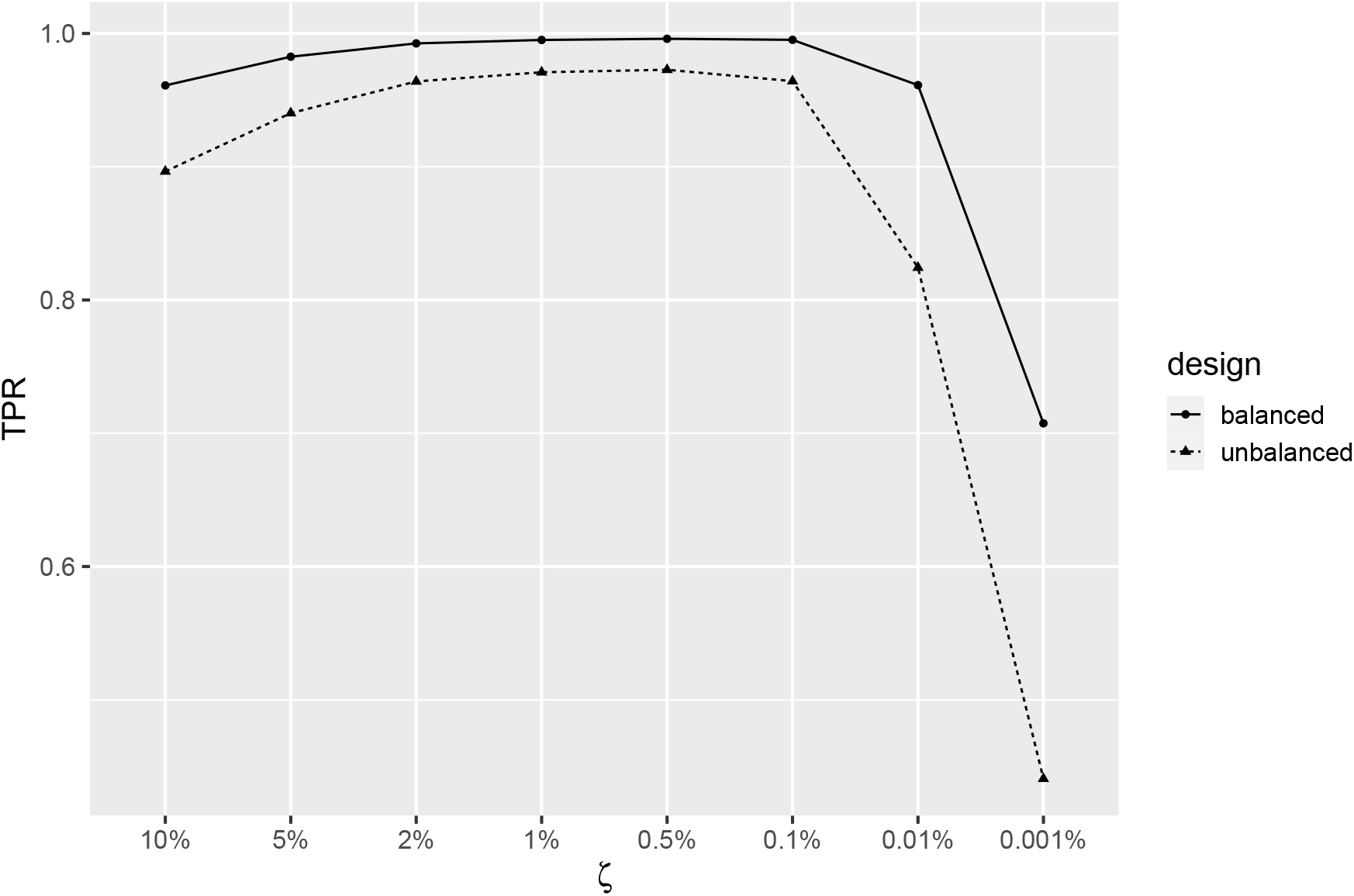
Plot of TPR for different choices of *ζ* for ComBat+Cor. The results were simulated based on the unbalanced/balanced group-batch design for the *bladderbatch* study.

**Figure 3:**
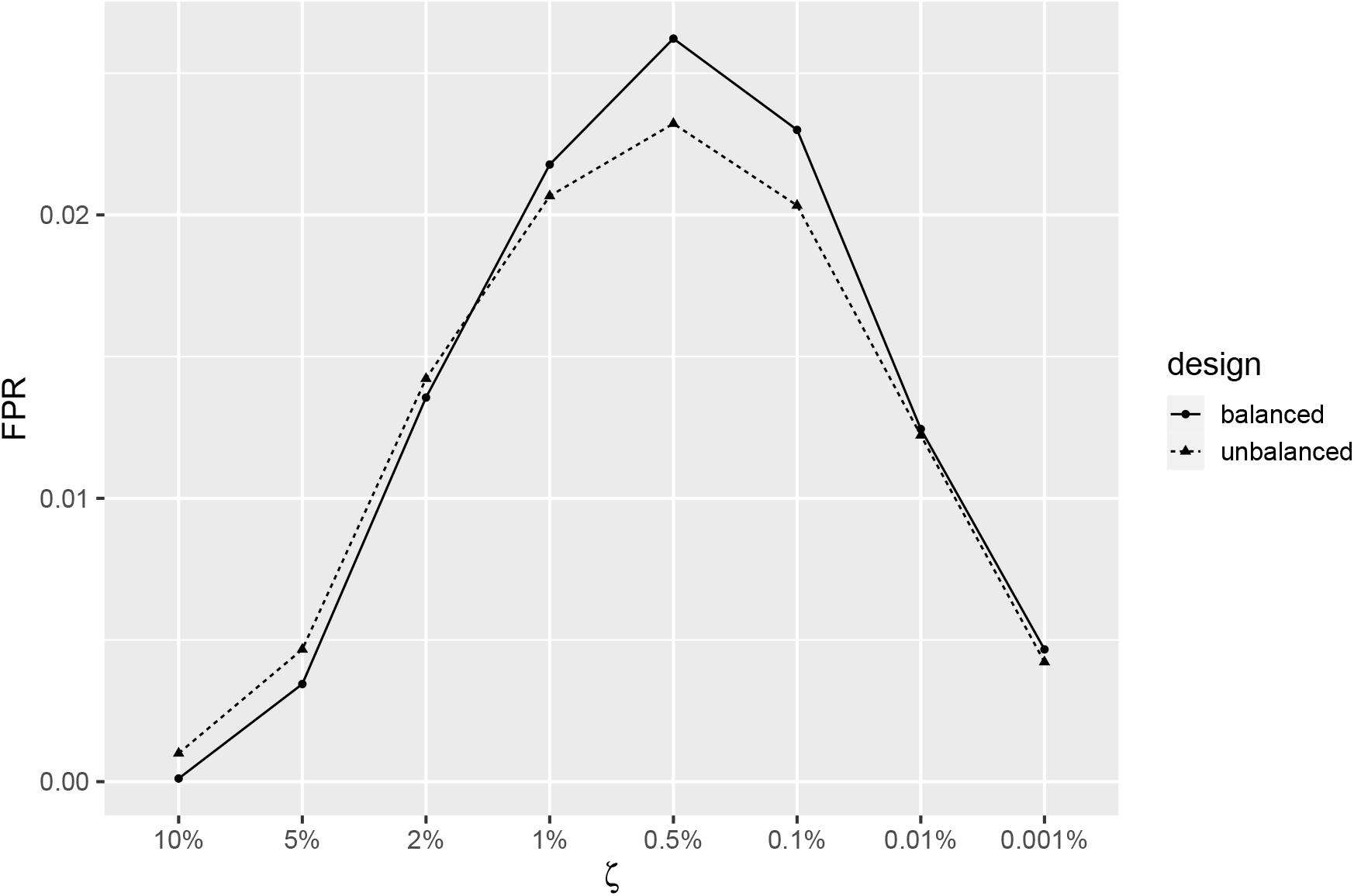
Plot of FPR for different choices of *ζ* for ComBat+Cor. The results were simulated based on the unbalanced/balanced group-batch design for the *bladderbatch* study.

Because the mean and variance batch effects were small in the original data, we conducted a second simulation with the *bladderbatch* data where we introduced large mean and variance batch effects to examine the performance of ComBat and ComBat+Cor (*ζ* = 1%) under more difficult conditions (see Table 2). We have two key observations: first, ComBat led to exaggerated significance in all cases where mean batch effects existed in an unbalanced design, and the level of exaggeration did not appear to have a relationship with the size of mean batch effects. Second, the size of variance batch effects had a strong impact on the performance of both ComBat and ComBat+Cor. When the variance batch effects were small, ComBat+Cor had much lower FPR than ComBat, which was clearly exaggerated in this case, in exchange for slightly worse TPR. This suggested that ComBat+Cor was a better choice than ComBat for small variance batch effects. When the variance batch effects were large, the TPR of ComBat+Cor reduced significantly while the exaggerated significance problem of ComBat disappeared, which suggested ComBat+Cor was overly conservative and less desirable than ComBat in this case. This is probably due to the advantage that ComBat has in dealing with variance batch effects and the fact that large variance batch effects would inflate the residual variance estimate and thus reduce statistical power.

For the comparison among all the approaches included in simulation, we found that for unbalanced design (Table 2), ComBat+Cor (*ζ* = 1%) consistently had FPR lower than 5% while maintaining a good statistical power. Notably, ComBat+Cor (*ζ* = 1%) was a better choice than the one-step approach as the one-step approach also tended to have exaggerated significance and decreased power when variance batch effects were large. The T-test without batch correction predictably performed the worst among all the candidates, which demonstrated the necessity for adjusting for batch effects in the data. When the group-batch design was balanced (Table 3) there were no significant differences among all approaches, except that the unadjusted T-test was still the worst performing approach. In general, we observed that ComBat+Cor was a safer choice than ComBat across all scenarios and protected against large FPR and consequent exaggerated significance for unbalanced group-batch designs.

**Table 3:** Results from *bladderbatch* simulation with balanced design. For each approach, results were obtained under the conditions where the mean and variance batch effects could be null (N), small (S) or large (L). For each condition, the results were formatted as FPR (TPR).

| Approach | Mean(N) |  |  | Mean(S) |  |  | Mean(L) |  |  |
| --- | --- | --- | --- | --- | --- | --- | --- | --- | --- |
|  | Var(N) | Var(S) | Var(L) | Var(N) | Var(S) | Var(L) | Var(N) | Var(S) | Var(L) |
| T-test | 5.0% | 5.3% | 4.9% | 0.9% | 0.9% | 1.9% | 0.0% | 0.0% | 0.2% |
|  | (99.9%) | (99.9%) | (97.9%) | (99.9%) | (99.9%) | (97.5%) | (98.0%) | (97.4%) | (93.7%) |
| Benchmark | 5.0% | 5.3% | 5.1% | 4.9% | 5.0% | 4.8% | 5.2% | 5.1% | 5.2% |
|  | (99.9%) | (99.9%) | (99.9%) | (99.9%) | (99.9%) | (100%) | (99.9%) | (100%) | (99.9%) |
| One-step | 5.0% | 5.2% | 4.6% | 4.9% | 4.8% | 4.6% | 5.2% | 4.9% | 4.8% |
|  | (99.9%) | (99.9%) | (97.8%) | (99.9%) | (99.9%) | (97.9%) | (99.9%) | (100%) | (97.6%) |
| ComBat | 5.2% | 6.9% | 0.0% | 4.6% | 6.8% | 0.1% | 5.4% | 6.9% | 0.0% |
|  | (99.9%) | (99.9%) | (99.4%) | (99.9%) | (99.9%) | (99.2%) | (99.9%) | (100%) | (98.8%) |
| ComBat+Cor($\zeta = 10\%$) | 0.0% | 0.1% | 0.0% | 0.0% | 0.0% | 0.0% | 0.0% | 0.0% | 0.0% |
|  | (98.5%) | (99.1%) | (91.2%) | (99.0%) | (99.0%) | (91.0%) | (99.0%) | (98.8%) | (89.3%) |
| ComBat+Cor($\zeta = 1\%$) | 2.6% | 3.9% | 0.0% | 2.3% | 4.0% | 0.0% | 2.8% | 4.0% | 0.0% |
|  | (99.8%) | (99.8%) | (99.0%) | (99.9%) | (99.9%) | (98.8%) | (99.9%) | (100%) | (98.4%) |
| ComBat+Cor($\zeta = 0.1\%$) | 1.2% | 3.6% | 0.0% | 1.1% | 6.2% | 0.1% | 2.1% | 6.4% | 0.0% |
|  | (99.3%) | (99.8%) | (98.9%) | (99.9%) | (99.9%) | (99.2%) | (99.9%) | (100%) | (98.7%) |
| ComBat+Cor($\zeta = 0.001\%$) | 0.0% | 0.0% | 0.0% | 0.0% | 1.7% | 0.0% | 0.0% | 2.5% | 0.0% |
|  | (41.5%) | (69.6%) | (47.1%) | (32.9%) | (99.9%) | (94.9%) | (54.2%) | (100%) | (96.6%) |

### 3.2 Example 2: Towfic et al. [2014]

Towfic et al. [2014] conducted an experiment to compare the effects of Copaxone and Glatimer, which are immunomodulators used to treat multiple sclerosis, and that was also used by Nygaard et al. [2016] to illustrate how ComBat can lead to exaggerated significance for an unbalanced batch-group design. There were 34 samples treated with Copaxone that were compared with 11 samples treated with Glatimer. In total, there were 17 batches and the batch-group design was highly unbalanced. Following the data processing and analysis procedure of Nygaard et al. [2016], there were 1,928 genes found to be significant at the 5% FDR threshold using ComBat adjusted data. We subsequently used ComBat+Cor (*ζ* = 1%) to adjust for correlations introduced by the unbalanced batch-group design, and found no genes were significant at 5% FDR level. However, we recognize that different models for differential expression, including mixed effects effects models, have led to deferentially expressed genes in this data set (Towfic et al., 2017, Nygaard et al., 2017). These can be further explored in the future with ComBat+Cor, but for the sake of this work, our goal was to recreate the work of Nygaard et al. [2016]. The simulation results based on this data (Figure 4(a)) uncovered that the statistical significance was highly exaggerated by ComBat and there was a strong need of adjustment for the unbalanced design, which is also consistent with the finding of Nygaard et al. [2016].

**Figure 4:**
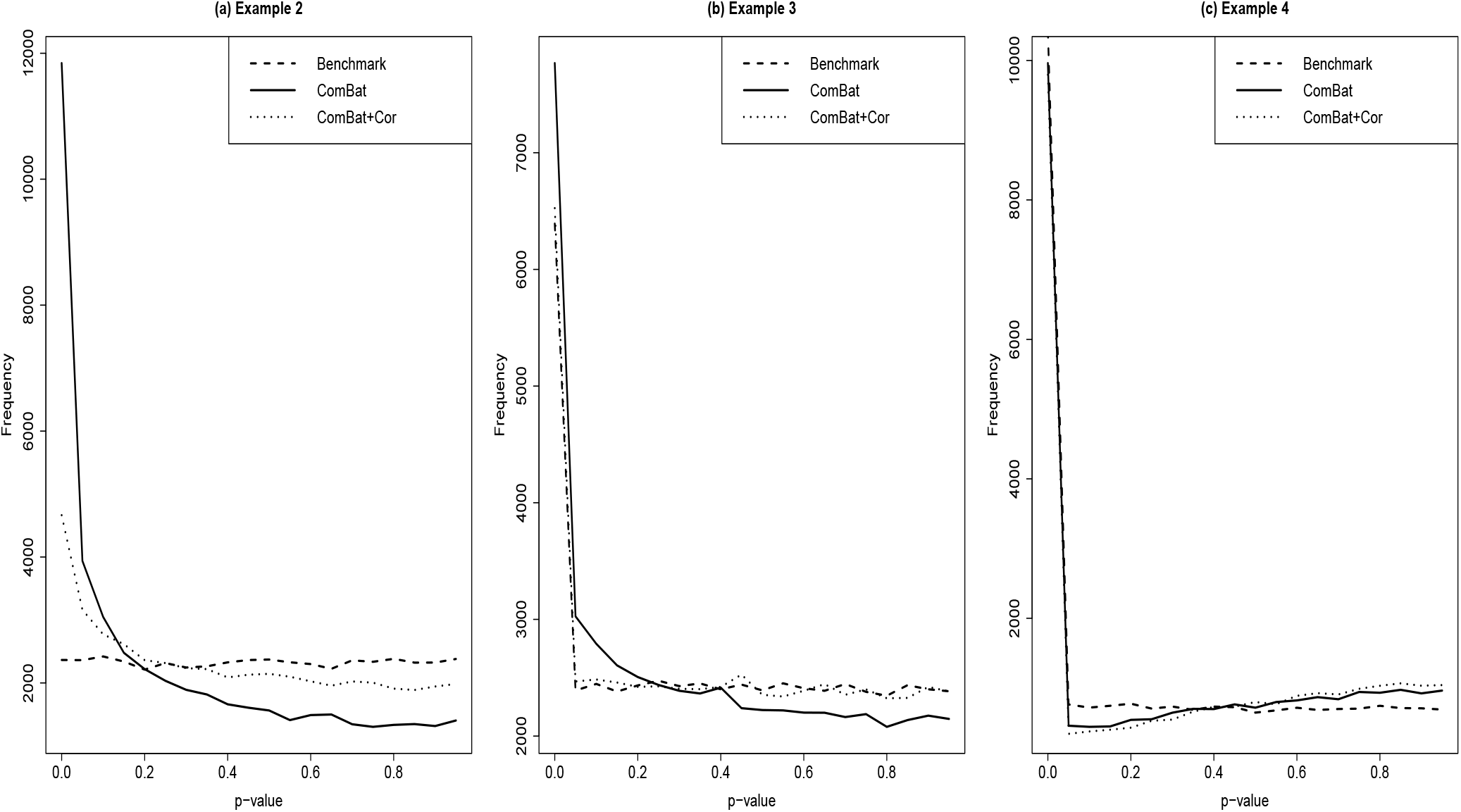
Simulation results for examples 2, 3 and 4. In each plot, we illustrate the distributions of the *p*-values for the benchmark approach, ComBat and ComBat+Cor. (a) Simulation results based on Towfic et al. [2014]. (b) Simulation results based on Johnson et al. [2007]. (c) Simulation results based on the TB data for comparing progressors versus non-progressors.

### 3.3 Example 3: Johnson et al. [2007]

Johnson et al. [2007] demonstrated ComBat using a dataset on the comparison of TAL1 inhibited cells. The experiment has 30 samples and 3 batches. The number of treated/control samples in each batch is batch 1: 6/2, batch 2: 3/4 and batch 3: 9/6. Importantly, Batch 3 consisted of technical replicates of the samples from Batches 1 and 2. Nygaard et al. [2016] also used this experiment to illustrate the exaggerated significance problem in ComBat. Following their processing procedure, we found 730 significant genes at 5% FDR using ComBat, but only 269 genes significant at 5% FDR using ComBat+Cor (*ζ* = 1%). Simulation results (Figure 4(b)) suggested that ComBat had a mild exaggerated significance problem due to the unbalanced design, which is consistent with the finding of Nygaard et al. [2016]. ComBat+Cor provides better control of the FDR and corrects the previously reported exaggerated significance problem.

### 3.4 Example 4: progressors versus non-progressors in tuberculosis

We present a final example of tuberculosis (TB) gene expression datasets which have been used to detect differentially expressed genes that distinguish progressors from non-progressors in TB (Zak et al., 2016, Suliman et al., 2018, Leong et al., 2018). This data example had three batches with each batch from a separate study. The ratios of the number of progressors and the number of non-progressors in each batch were 77/104, 95/304, and 0/19. We chose *ζ* as 0.1% as guided by the recommended range and the sample size, and simulation results supported this choice (Figure 5). Of 24,391 genes, We found 9,659 significant genes at 5% FDR using ComBat and 8,403 significant genes at 5% FDR using ComBat+Cor (*ζ* = 0.1%). We observed that the significant genes found by ComBat contained all the genes found by ComBat+Cor. Our simulation results (Figure 4(c)) showed that most of the discovered genes were expected to be differentially expressed, as both ComBat and ComBat+Cor had diminished significance in the simulated data due to large variance batch effects.

**Figure 5:**
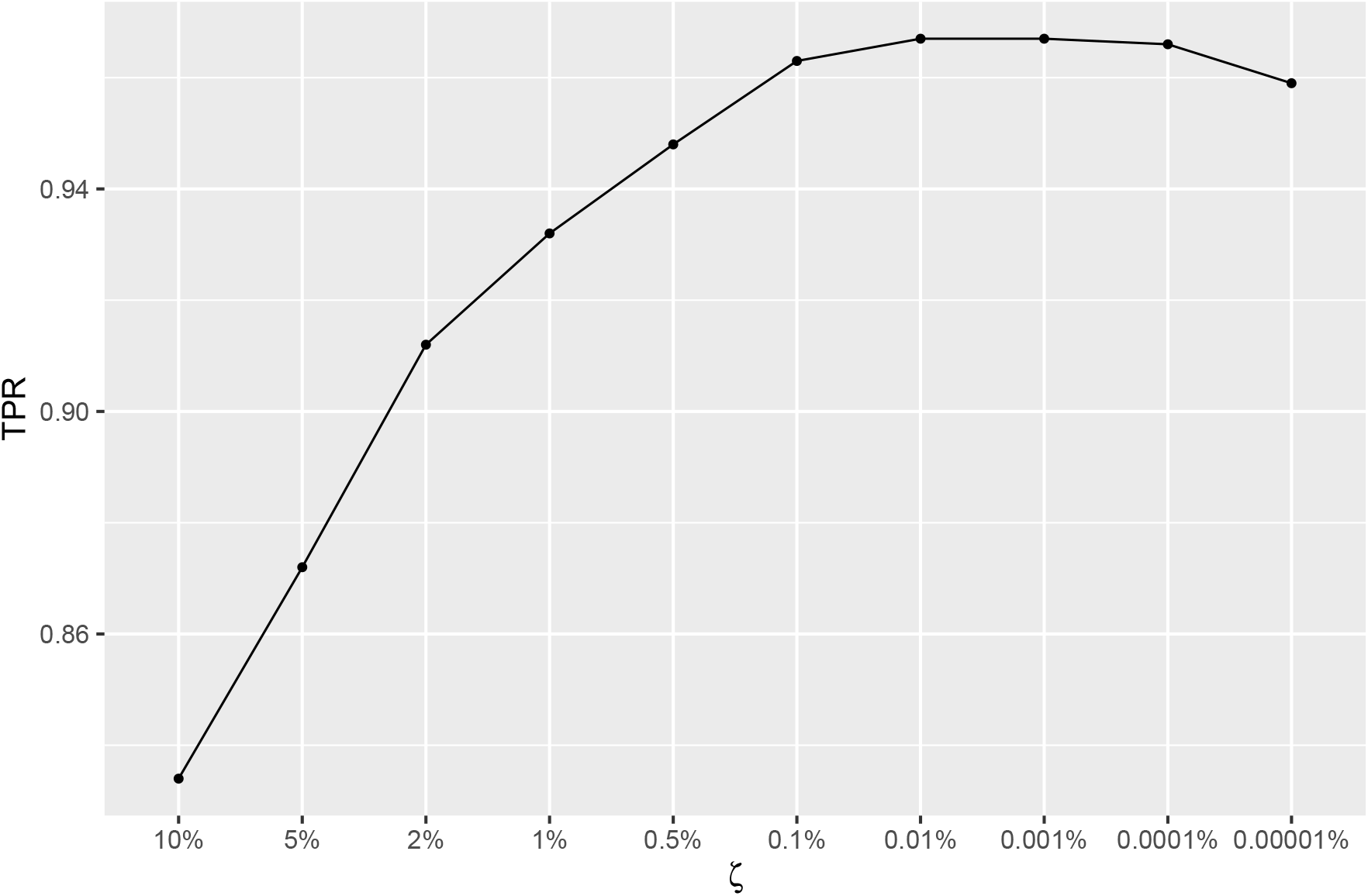
Plot of TPR for different choices of *ζ* for ComBat+Cor. The results were simulated based on the original TB dataset.

## 4 Discussion

ComBat is an established tool for batch effect adjustment, but we have shown it can often lead to inflated (or deflated) significance in gene expression studies, particularly for unbalanced group-batch designs. The exaggerated significance of ComBat results from the fact that samples are correlated after batch adjustment, because removing the estimated mean batch effect from the original data relies on all observations within a batch. To avoid this problem, downstream methods must account for the correlation induced by batch adjustment.

We have shown that the sample correlation matrix can be derived based on the group-batch design and should be incorporated into downstream analyses. Because the derived sample correlation matrix is not full-rank, we proposed a procedure that adds a small amount of random noise into the data using a parameter *ζ*. This recovers approximate estimability of the covariance structure and enables approximation through a spectral decomposition approach. The ComBat three-step approach with a correlation adjustment, ComBat+Cor, is defined as follows: (1) use the original ComBat to obtain batch-corrected data; (2) compute the approximated sample correlation matrix; (3) conduct downstream modeling with appropriate accommodations for correlated data, such as GLS, using the estimated sample correlation matrix.

Our simulation results based on a real dataset with substantial group-batch imbalance and both mean and variance batch effects demonstrate that accounting for the sample correlation matrix via Combat+Cor provides consistent control of the false positive rate for differential expression analysis. This is especially important considering the exaggerated significance problem of ComBat in unbalanced group-batch designs. ComBat+Cor is consistently more conservative than ComBat regardless of the choice of *ζ* and thus protects against inflated FDR. For a recommended choice of *ζ* (i.e., between 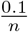 and 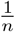), ComBat+Cor can also achieve good statistical power, making it more desirable than ComBat for unbalanced group-batch designs without large variance batch effects. It is also noteworthy that ComBat+Cor maintains a better balance between TPR and FPR and is more flexible than the one-step approach for unbalanced group-batch designs, as the one-step approach is not always available. We still recommend using ComBat for balanced group-batch designs, as it consistently yields higher statistical power and has no signs of exaggerated significance in the balanced designs.

We caution readers that ComBat+Cor is less desirable for data with large variance batch effects, as it may become too conservative and underreport the number of truly significant features. The exaggerated significance problem for ComBat may not always be present in data examples where batches have large variance batch effects. Given ComBat+Cor actually loses TPR in exchange for a reduction in FPR, such a tradeoff would be undesirable when the ComBat approach does not lead to exaggerated significance. Therefore, we recommend using ComBat for data with large variance batch effects and ComBat+Cor for data with small variance batch effects. To facilitate the decision-making process, we illustrate the guidance about the choice of ComBat and ComBat+Cor in Figure 6.

**Figure 6:**
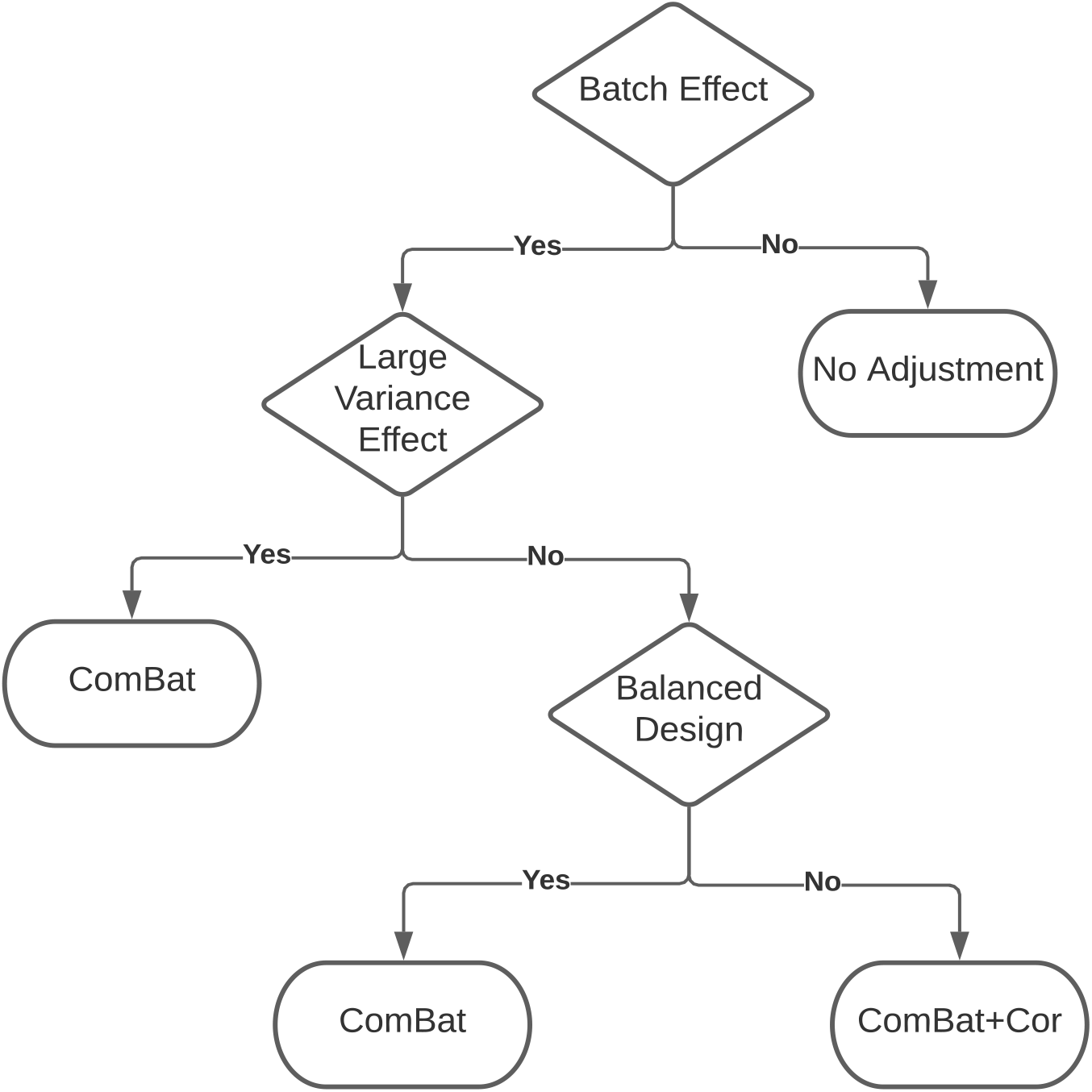
Guidance about the choice of ComBat and ComBat+Cor for addressing the exaggerated significance problem in batch correction.

Future research is needed in the following three directions: First, more in-depth discussions about the role of variance batch effects in downstream analyses as well as the inference of biological effects are needed. Second, correlations in the batch-corrected data given by ComBat may be partially due to empirical Bayes (EB) processing, and therefore learning the impact of EB processing is necessary for a comprehensive understanding of the sample correlations induced by ComBat. Third, customary solutions are needed for other popular methods such as DESeq (Anders and Huber, 2010, Love et al., 2014), limma (Smyth, 2005) or edgeR (Robinson et al., 2010), where sample correlations and exaggerated significance may exist after batch correction.

## 5 Software and Code

Software in the form of R code, together with a sample input data set and complete documentation is available online at GitHub (https://github.com/tenglongli/ComBatCorr). Function for outputting the adjusted *p*-values or the 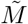 matrix (function name: design adj) is also available in GitHub (https://github.com/jtleek/sva-devel) and will be available in future versions of the sva package in Bioconductor (https://bioconductor.org/packages/release/bioc/html/sva.html).

## 6 Supplementary Material

Supplementary material is available online at http://biorxiv.org.

## Acknowledgments

This work was supported by funds from the NIH: 5U01CA220413 and 5R01GM127430.

