## Supplementary Material for "Overcoming the impacts of two-step batch effect correction on gene expression estimation and inference"

#### 1 Derivation of the sample covariance matrix for the two-step approach

Based on the Frisch-Waugh-Lovell theorem (Frisch and Waugh, 1933, Lovell, 1963, 2008), the estimate for batch effect $\hat{\beta}_{2g}$ in the regression model $Y_g = X_1\beta_{1g} + X_2\beta_{2g} + \epsilon_g, \epsilon_g \sim N(0, \sigma_g^2 I)$, is the same as the $\hat{\beta}_{2g}$ in the following regression model:

$$PY_g = PX_2\beta_{2g} + \epsilon_g, \epsilon_g \sim N(0, \sigma_g^2 I) \quad (1)$$

where $P = I - X_1(X_1^T X_1)^{-1}X_1^T$. We will also have $\hat{\beta}_{2g} = (X_2^T P X_2)^{-1} X_2^T P Y_g$.

Model (1) the first step of the two-step batch adjustment. In the second step of the two-step batch adjustment, the batch adjusted data $\tilde{Y}_g$ is obtained as:

$$\tilde{Y}_g = Y_g - X_2\hat{\beta}_{2g} = (I - X_2(X_2^T P X_2)^{-1} X_2^T P)Y_g = (I - H_{12})Y_g \quad (2)$$

*To whom correspondence should be addressed.

where $H_{12}$ is:

$$H_{12} = X_2(X_2^T P X_2)^{-1} X_2^T P = X_2(X_2^T(I - X_1(X_1^T X_1)^{-1} X_1^T)X_2)^{-1} X_2^T(I - X_1(X_1^T X_1)^{-1} X_1^T) \quad (3)$$

Therefore, the batch adjusted data has covariance matrix $\sigma_g^2(I - H_{12})(I - H_{12})^T$.

It is noteworthy that $X_1$ should also include the all-ones vector $\mathbf{1}$ when there is an reference batch. That is $H_{12} = X_2(X_2^T(I - X_0(X_0^T X_0)^{-1} X_0^T)X_2)^{-1} X_2^T(I - X_0(X_0^T X_0)^{-1} X_0^T)$ where $X_0 = [\mathbf{1}, X_1]$.

### 2 The relationship between biological effect estimates and batch design

In this section, we will show the relationship between biological effect estimates and batch design. Without loss of generality, the regression model (for each gene) is formulated as $Y = \alpha + X_1\beta_1 + X_2\beta_2 + \epsilon, \epsilon \sim N(0, \sigma^2 I)$, where $\alpha$ is the background gene expression. $X_1$ represents the biological groups (assume there are two biological groups) and $X_2$ represents the batch design. Furthermore, we define the matrix $X = [X_2, X_1]$ and the matrix $V = [\mathbf{1}, X]$. We also define sample variance $\hat{\sigma}_{xx}$ and covariances $\hat{\sigma}_{xy}$ as follows:

$$\hat{\sigma}_{xx} = \frac{1}{n}\sum_{i=1}^{n}(x_i - \bar{x})^2 \quad (4)$$

$$\hat{\sigma}_{xy} = \frac{1}{n}\sum_{i=1}^{n}(x_i - \bar{x})(y_i - \bar{y}) \quad (5)$$

Our goal is to derive the least square estimate $\hat{\beta}_1$ and its variance based on the sample variance-covariance matrix of $X$. It's known that the least square estimate has the matrix form $(V^T V)^{-1} V^T Y$. Specifically, $V^T V$ is the following block matrix:

$$V^T V = \begin{pmatrix} n & n\bar{X}^T \\ n\bar{X} & X^T X \end{pmatrix} \quad (6)$$

where $\bar{X} = [\bar{X}_2, \bar{X}_1]^T$.

The inverse of $V^T V$ then becomes:

$$(V^T V)^{-1} = \begin{pmatrix} n^{-1} + n^{-1}\bar{X}^T S_{XX}^{-1}\bar{X} & -n^{-1}\bar{X}^T S_{XX}^{-1} \\ -n^{-1}S_{XX}^{-1}\bar{X} & n^{-1}S_{XX}^{-1} \end{pmatrix} \quad (7)$$

Furthermore, the covariance matrix $S_{XX}$ is the following block matrix:

$$S_{XX} = \begin{pmatrix} S_{22} & S_{21} \\ \\ S_{12} & \hat{\sigma}_{11} \end{pmatrix} \tag{8}$$

where $S_{22}$ is the covariance matrix of $X_2$, $S_{21}$ is the covariance matrix between $X_2$ and $X_1$ ($b$ rows and 1 column; $b$ is the number of batch indicators), and $S_{12}$ is just the transpose of $S_{21}$. The inverse of $S_{XX}$ is:

$$S_{XX}^{-1} = \begin{pmatrix} S_{22}^{-1} + S_{22}^{-1}S_{21}(\hat{\sigma}_{11} - S_{12}S_{22}^{-1}S_{21})^{-1}S_{12}S_{22}^{-1} & -S_{22}^{-1}S_{21}(\hat{\sigma}_{11} - S_{12}S_{22}^{-1}S_{21})^{-1} \\ \\ -(\hat{\sigma}_{11} - S_{12}S_{22}^{-1}S_{21})^{-1}S_{12}S_{22}^{-1} & (\hat{\sigma}_{11} - S_{12}S_{22}^{-1}S_{21})^{-1} \end{pmatrix} \tag{9}$$

Plugging in the above expression of $S_{XX}^{-1}$ into the block matrix in (7) will give the complete form of $(V^TV)^{-1}$, whose elements are all sample means or sample variances/covariances. To isolate $\hat{\beta}_1$, only the last row of $(V^TV)^{-1}$ is needed:

$$(V^TV)^{-1}_{(b+2)1} = n^{-1}[(\hat{\sigma}_{11} - S_{12}S_{22}^{-1}S_{21})^{-1}S_{12}S_{22}^{-1}\bar{X}_2 - \bar{X}_1(\hat{\sigma}_{11} - S_{12}S_{22}^{-1}S_{21})^{-1}] \tag{10}$$

$$[(V^TV)^{-1}_{(b+2)2}, \ldots, (V^TV)^{-1}_{(b+2)(b+1)}] = -n^{-1}(\hat{\sigma}_{11} - S_{12}S_{22}^{-1}S_{21})^{-1}S_{12}S_{22}^{-1} \tag{11}$$

$$(V^TV)^{-1}_{(b+2)(b+2)} = n^{-1}(\hat{\sigma}_{11} - S_{12}S_{22}^{-1}S_{21})^{-1} \tag{12}$$

$\hat{\beta}_1$ is straightforward given the following expression of $V^TY$:

$$V^TY = \begin{pmatrix} n\bar{Y} \\ \\ nS_{2Y} + n\bar{Y}\bar{X}_2 \\ \\ n\hat{\sigma}_{1Y} + n\bar{X}_1\bar{Y} \end{pmatrix} \tag{13}$$

The expression of $\hat{\beta}_1$ is then:

$$\hat{\beta}_1 = \frac{\hat{\sigma}_{1Y} - S_{12}S_{22}^{-1}S_{2Y}}{\hat{\sigma}_{11} - S_{12}S_{22}^{-1}S_{21}} \tag{14}$$

and its variance is:

$$\text{Var}(\hat{\beta}_1) = \frac{\hat{\sigma}^2}{n}(\hat{\sigma}_{11} - S_{12}S_{22}^{-1}S_{21})^{-1} \tag{15}$$

where $\hat{\sigma}^2$ is the estimated residual variance in regression.

### 3 The relationship between $H_{12}$ and $S_{12}$

Without loss of generality, we assume there is a reference batch and thus $H_{12} = X_2(X_2^T(I - X_0(X_0^TX_0)^{-1}X_0^T)X_2)^{-1}X_2^T(I - X_0(X_0^TX_0)^{-1}X_0^T)$ where $X_0 = [\mathbf{1}, X_1]$. Following the regression framework in the previous section, we can derive the expression of $(X_0^T X_0)^{-1} X_0^T X_2$. First, we have:

$$(X_0^T X_0)^{-1} = \begin{pmatrix} n^{-1} + n^{-1}\bar{X}_1^2 \hat{\sigma}_{11}^{-1} & -n^{-1}\bar{X}_1 \hat{\sigma}_{11}^{-1} \\ -n^{-1}\bar{X}_1 \hat{\sigma}_{11}^{-1} & n^{-1}\hat{\sigma}_{11}^{-1} \end{pmatrix} \tag{16}$$

and:

$$X_0^T X_2 = \begin{pmatrix} n\bar{X}_2 \\ nS_{12} + n\bar{X}_1 \bar{X}_2 \end{pmatrix} \tag{17}$$

where $\bar{X}_2 = [\bar{X}_{21}, \bar{X}_{22}, \ldots, \bar{X}_{2b}]$, a 1 by $b$ vector whose elements are means of the batch indicators in $X_2$.

Taken together, we have the expression of $(X_0^T X_0)^{-1} X_0^T X_2$ as follows:

$$(X_0^T X_0)^{-1} X_0^T X_2 = \begin{pmatrix} \bar{X}_2 - \bar{X}_1 \hat{\sigma}_{11}^{-1} S_{12} \\ \hat{\sigma}_{11}^{-1} S_{12} \end{pmatrix}_{2\times b} \tag{18}$$

#### 3.1 Special case: balanced designs

For a balanced group-batch design, the elements in $S_{12}$ are all 0, which means:

$$(X_0^T X_0)^{-1} X_0^T X_2 = \begin{pmatrix} \bar{X}_2 \\ \mathbf{0} \end{pmatrix}_{2\times b} \tag{19}$$

Based on (19), we can derive the expression of $(I - X_0(X_0^T X_0)^{-1} X_0^T)X_2$ as follows:

$$(I - X_0(X_0^T X_0)^{-1} X_0^T)X_2 = \begin{pmatrix} X_{211} - \bar{X}_{21} & X_{212} - \bar{X}_{22} & \cdots & X_{21b} - \bar{X}_{2b} \\ X_{221} - \bar{X}_{21} & X_{222} - \bar{X}_{22} & \cdots & X_{22b} - \bar{X}_{2b} \\ \vdots & \vdots & \cdots & \vdots \\ X_{2n1} - \bar{X}_{21} & X_{2n2} - \bar{X}_{22} & \cdots & X_{2nb} - \bar{X}_{2b} \end{pmatrix}_{n\times b} \tag{20}$$

It is clear that (20) is just the centered version of $X_2$ (denoted as $X_2^c$), and so we can express $H_{12}$ as:

$$H_{12} = X_2(X_2^T X_2^c)^{-1}(X_2^c)^T \tag{21}$$

With $H_{12}$ in the form of (21), the correlation between two samples from two different batches in the matrix $M$ is $\frac{1}{n_r}$ where $n_r$ is the sample size of the reference batch. If a reference batch is not used, as is the case with most applications of ComBat, the correlations are not as straightforward, but can be derived using a similar procedure, and lead to the same conclusion (covariance only depends on the batch design).

To summarize, when the group-batch design is balanced, $H_{12}$ is solely a function of the batch design $X_2$ and has nothing to do with the group design $X_1$. Therefore, removing batch effects in the first step won't result in the *endogeneity* issue for the second step. When the group-batch design is unbalanced, $H_{12}$ depends on both the batch design $X_2$ and the group design $X_1$. The relationship between $H_{12}$ and $S_{12}$ (and thus $X_1$) can be derived by plugging the expression (18) into the expression of $H_{12}$, which will not be detailed here. Most importantly, when the group-batch design is unbalanced, the covariance vector $S_{12}$ will not be $\mathbf{0}$ and thus $H_{12}$ will depend on $X_1$ and the strength of such dependence is characterized by $S_{12}$.
